# Taxonomic investigation of the zooplanktivorous Lake Malawi cichlids *Copadichromis mloto* (Iles) and *C. virginalis* (Iles)

**DOI:** 10.1101/2022.03.10.483679

**Authors:** G.F. Turner, D.A. Crampton, A. Hooft van Huysduynen, H. Svardal

**Affiliations:** School of Natural Sciences, Bangor University, Bangor, Gwynedd LL57 2UW, UK; Vertebrates Division, Natural History Museum, Cromwell Road, London SW7; University of Antwerp, Universiteitsplein 1, 2610 Wilrijk, Belgium; Naturalis Biodiversity Center, 2300 RA Leiden, The Netherlands

**Keywords:** cichlid fish, Lake Malawi, utaka, systematics, whole genome sequencing, geometric morphometrics

## Abstract

The taxonomic status of the zooplanktivorous cichlids *Copadichromis mloto* and *C. virginalis* has been confused since their original descriptions by lles in 1960. While two forms of *C. virginalis*, Kaduna and Kajose, were distinguished in the type material, *C. mloto* has not been positively identified since its original description. Here we re-examined the types as well as 54 recently collected specimens from multiple sampling locations. Genome sequencing of 51 recent specimens revealed two closely related but reciprocally monophyletic clades. Geometric morphological analysis indicated that one clade morphologically encompasses the type specimens of *C. virginalis* identified by Iles as the ‘Kaduna’ form, including the holotype, while the other clade encompasses not only the paratypes identified as the ‘Kajose’ form, but also the type series of *C. mloto*. Given that all three forms have the same type locality, that there are no meristic or character states to differentiate them and that there are no records of adult male *C. mloto* in breeding colours, we conclude that the ‘Kajose’ form previously identified as *C. virginalis* represents relatively deeper bodied sexually active or maturing individuals of *C. mloto*.

## Introduction

The Lake Malawi haplochromine cichlids represent the most species-rich vertebrate adaptive radiation known, comprising around 800 species (Konings, 2016) rapidly evolved from a common ancestor in a single lake (Malinsky et al., 2018; Svardal et al., 2020), and as such they represent a particularly difficult taxonomic challenge (Snoeks, 2004). The utaka are a group of zooplankton feeding cichlids that are both ecologically and commercially important as a food fish (Turner, 1996). Most utaka species are currently assigned to the genus *Copadichromis*, characterised by their relatively small, highly protrusible mouths and numerous long gill rakers (Eccles &Trewavas, 1989; Konings, 2016). They are generally silvery, countershaded and carry several dark spots on their flanks, although this is obscured in the reproductively active males that are conspicuously dark blue or black (Konings, 2016). A few species, known as ‘pure utaka’ lack flank spots entirely (Iles, 1960). The status of these has been confused for over 60 years, since the descriptions of *Copadichromis mloto* (Iles, 1960) and *Copadichromis virginalis* (Iles, 1960), both described from material collected at Nkhata Bay in the middle of the western shore of the lake. The former species was distinguished from the latter by its more slender build, but all other morphometric traits and all meristic counts overlapped (Iles, 1960). While the original descriptions presented information on male breeding characteristics (often a key feature for discriminating closely-related cichlid species) for *C. virginalis*, none were given for *C. mloto* which all appeared to be spent or reproductively inactive individuals. In the intervening years, no breeding adults of *C. mloto* were positively identified in collections, although Konings (2016) illustrated specimens proposed to be *C. mloto* that resemble those commonly assigned to *C. virginalis* in commercial trawl catches (Turner, 1996), but which had been assigned as *C. mloto* in earlier publications (e.g. Axelrod &Burgess, 1986). Further confusing matters, Iles’s original description of *C. virginalis* discussed two distinct sympatric forms which he recorded local fishermen referring to as ‘Kaduna’ (including the holotype) and ‘Kajose’ (included in the type series). Subsequent authors (e.g. Turner, 1996; Konings, 2016) suggested that these may be different species – a possibility tentatively discussed by Iles. In the course of a wider investigation of the Lake Malawi cichlid fauna, we were able to obtain a large number of specimens of ‘pure utaka’ from a number of locations in 2016-17, which we used to investigate the status of *C. mloto* and *C. virginalis* using a phylogeny constructed from whole genome sequences, coupled with geometric morphometric comparisons with Iles’ type material.

## Methods and Material

This study was based on the type material of *Haplochromis mloto* and *H. virginalis* examined and photographed in the Natural History Museum in London, along with 54 specimens collected from various locations around Lake Malawi in 2016-2017. The types of *H. virginalis* were classified by Iles into Kaduna and Kajose forms: these were separately catalogued: the holotype is a female Kaduna morph. During our examinations, external morphology suggested that two of the specimens were misclassified (perhaps through a mix-up by a later researcher), with the 88.3mm SL individual in jar labelled as Kaduna looking more like a Kajose and the 96.5mm SL Kajose looking more like a Kaduna. We used the re-classified identities in our analyses.

The freshly collected specimens were purchased from fishermen, and if not already dead, euthanised with anaesthetic overdose (clove oil); the right pectoral fin was cut off and placed in a vial of pure ethanol; the specimen pinned, labelled and photographed before being preserved in formalin after rigor mortis had set in, before being washed and preserved in ethanol. Morphometric analysis was based on digital analysis of the field photographs, along photographs taken of the type material. Previous studies had examined meristics and morphological character states and did not find any diagnostic differences between the species (Iles, 1960; Eccles &Trewavas, 1989). Iles (1960) suggested that *C. mloto* had smaller teeth than similar species, but this was not supported by preliminary investigations – across species, large adult males were found to have strong simple teeth and smaller fish have relatively smaller teeth, generally bicuspid in the outer row and tricuspid in the inner rows. This was not investigated further.

For geometric morphometric (GM) analysis, tpsUTIL (Rohlf, 2004) was used to build a file from scaled photographs, co-ordinates were recorded by tpsDig2 ver 2 (Rohlf, 2015) using the landmark tool, and provided with scale factors. There were 15 homologous landmarks on the full specimen (Fig. 1). A CVS file containing the x and y coordinates of the landmarks for each specimen was then created and imported into MorphoJ (Klingenberg, 2011). Before Principal Components Analysis (PCA) was run on the geometric morphometrics data, a generalized Procrustes analysis (GPA) was applied on the landmark data, to mathematically remove non-shape variation (Rohlf &Slice, 1990; Bookstein, 1989; Parés-Casanova et al., 2020). This eliminated any morphological variations resulting from the size, position or orientation of the specimens. Then, a covariance matrix was generated from the resulting Procrustes shape coordinates and lastly the PCA was carried out. After this, transformation grids were created to visualise the landmark configurations of the shapes. Each point represents the consensus shape (mean shape of the sample), and the line represents the shape change that corresponds to an increase of 0.1 units of Procrustes distance in the direction of the PC1. The resulting PC scores and centroid sizes were then imported into IBM SPSS Statistics 27, and One-Way Analysis of Variance used to test for group differences among component scores and centroid sizes (CS: a measure of overall body size) among the following 5 groups: types of *H. mloto*, types of *H. virginalis* (Kaduna), types of *H. virginalis* ‘Kajose’, sequenced *C. mloto* and sequenced *C. virginalis*. Posthoc tests (Tukey) were used to identify significant differences among groups, after simultaneous Bonferroni correction. Correlations between CS and PC scores were also calculated to aid interpretation.

**Fig. 1.**
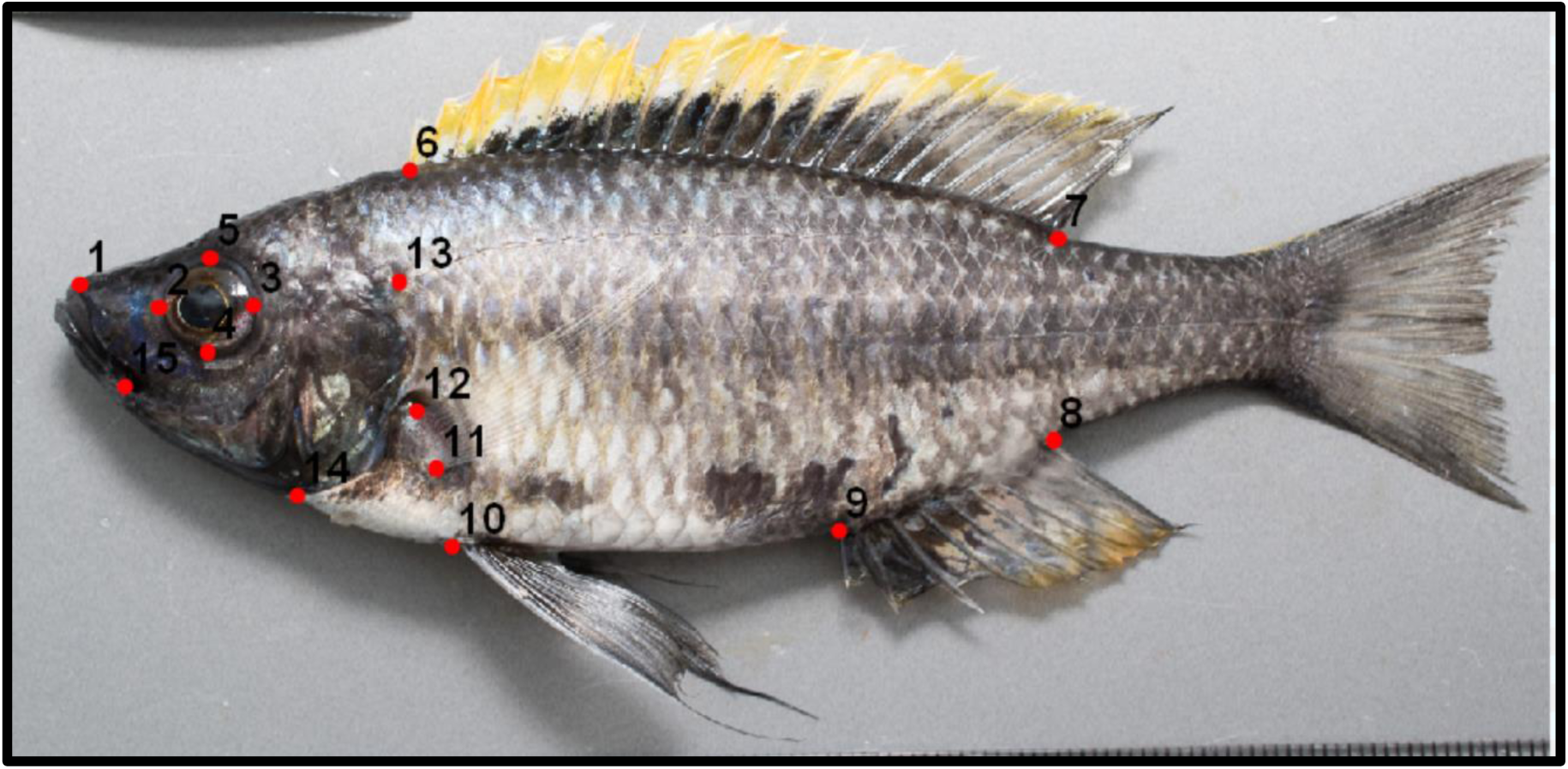
Landmarks used in Geometric Morphometric analysis of external body proportions (Specimen D12I04, *C. mloto*). (1) anterior tip of upper surface of maxilla (2) posterior reach of the eye (3) anterior reach of the eye (4) ventral reach of the eye (5) dorsal reach of the eye (6) anterior insertion point of dorsal fin (7) posterior insertion point of dorsal fin (8) posterior insertion point of anal fin (9) anterior insertion of anal fin (10) anterior/dorsal insertion of pelvic fin (11) anterior/ ventral insertion of pectoral fin (12) dorsal insertion of pectoral fin (13) point lateral line meets operculum (14) ventral-posterior extreme of preoperculum (15) anterior reach of the premaxillary groove.

DNA libraries for the above specimens collected in 2016-2017 as well as a single individual of the outgroup *Copadichromis chrysonotus* (Boulenger, 1908) were created and whole-genome sequenced on an Illumina HiSeq platform to individual coverages between 13.8-fold and 42.1-fold (median of 17.1 -fold). For this study reads were aligned to the *Astatotilapia calliptera* reference genome fAstCal1.2 (GCA_900246225.3 https://www.ncbi.nlm.nih.gov/assembly/GCF_900246225.1) using BWA-MEM (BWA version 0.7.17). The variant site detection was performed with bcftools (version 1.14, Danecek, et al., 2021). Any sites where more than 10% of mapped reads had a mapping quality zero, or where the overall mapping quality was less than 50 were masked. Sites for which mapping quality was significantly different between the forward and reverse strand (P<0.001) were also masked, as well as sites where the sum of overall depth for all samples was unusually high (>97.5 percentile) or low (<2.5 percentile). Heterozygous sites for which a binomial test showed a significantly biased read depth of the reference and alternative alleles (PHRED score >20) were also masked, representing between 0.02% and 0.15% of heterozygous sites per individual. Additionally, filters were applied to sites with excess heterozygosity (InbreedingCoeff < 0.2) and sites with >20% missing genotypes. Only single nucleotide polymorphisms (SNPs) were retained for genetic investigation. After filtering, 17,307,336 biallelic SNPs across 22 chromosomes remained. Three specimens, D20E07, D20F05, and D20F06 collected at Lake Malombe were excluded from genetic analysis because of possible cross-contamination in the sequencing data apparent from substantial excess heterozygosity.

Pairwise sequence differences were calculated in 100kb windows and averaged over the two haplotypes of each individual with a custom script based on scikit allel (available at https://github.com/feilchenfeldt/pypopgen3). To reconstruct genome-wide phylogenetic relationships, we summed pairwise genetic differences across all windows and constructed a neighbour-joining (NJ) tree using the Biophyton 1.79 Phylo package. We note that NJ has been shown to be a statistically consistent and accurate species tree estimator under incomplete lineage sorting (Rusinko &McPartlon, 2017), which is known to be pervasive among Malawi cichlid lineages (Malinsky et al., 2018). The resulting NJ tree was rooted using the *C. chrysonotus* sample as an outgroup. We implemented a block bootstrap by resampling with replacement the distance matrices of 100kb windows 1000 times, each sample of a size corresponding to the total number of matrices (8,519), computed an NJ tree for each of these samples, and tested in what percentage of the samples the topology of a given node of the original tree was supported. Finally, we also constructed a “split-tree” of samples where for each pairwise comparison the average within-individual heterozygosity (i.e. the pairwise sequence difference between the two haplotypes for each individual) was subtracted from the between-individual pairwise differences before constructing an NJ tree. We used the resulting NJ tree to obtain and estimate of the split time between *C. virginalis* and *C. mloto*, by averaging the distance of present-day samples to the node separating the two species, using the previously inferred Malawi cichlid mutation rate of 3.5 × 10^−9^ (confidence intervals: 1.6-4.7 × 10^−9^; Malinsky et al., 2018), the accessibility mask from our variant calling, and assuming a generation time of three years. We note that this estimate of split time assumes constant effective population sizes and mutation rates.

### Specimens examined

A representative selection of specimens is shown on Figure 2.

**Fig. 2.**
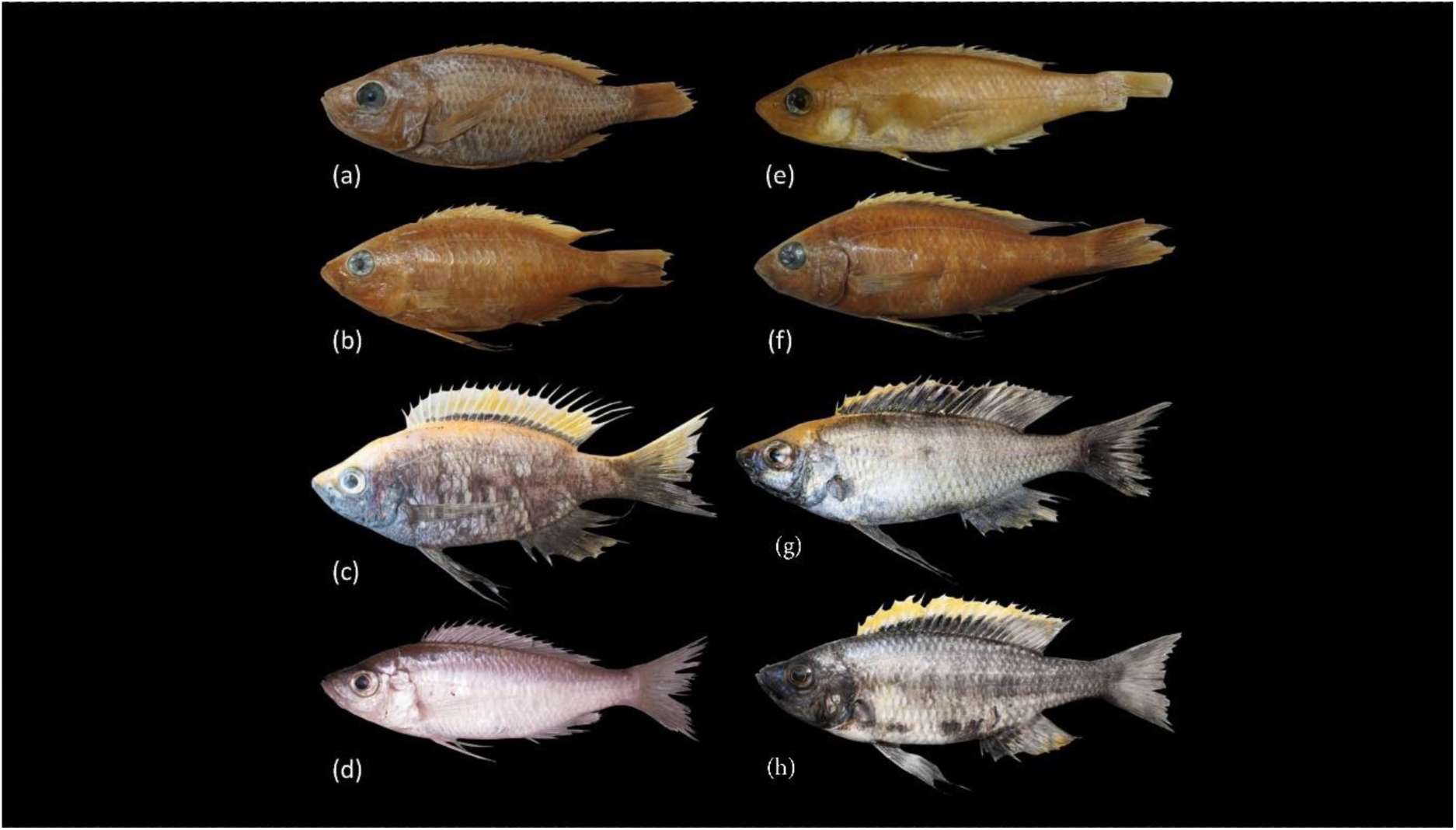
*Copdichromis virginalis*: (a) NHM 1962.10.18.21, holotype of *H. virginalis*, female 79.8 mm SL; (b) NHM 1962.10.18.22-30, paratype of *H. virginalis* (Kaduna morph), male; (c) D02G09, male, Nkhata Bay; *Copadichromis mloto*: (d) D17A03, female, Palm Beach, SE Arm; (e) NHM 1961.10.18.31, holotype of *H. mloto*: apparent female 92.4 mm SL; NHM 1961.10.18.40-50; (f) paratype of *H. virginalis* (Kajose morph), male; (g) D06E04, male, Chiweta Beach, Chilumba; (h) D12I04, male, trawled from 30-40m depth, SW of Makanjila.

#### *Copadichromis mloto* (Iles, 1960)

NHM 1962.10.18.21 Holotype of *Haplochromis mloto*, coll. T.D.Iles, Nkhata Bay, Lake Nyasa 79.8 mm SL NHM 1962.10.18.22-30 Paratypes of *Haplochromis mloto*, coll. T.D.Iles, Nkhata Bay, Lake Nyasa 9 specimens 76.8-114.7 mm SL

NHM 1961.10.18.40-50 Paratypes of *Haplochromis virginalis* (Kajose form), coll. T.D.Iles, Nkhata Bay, Lake Nyasa 10 specimens 70.4-111.2 mm SL

NHM 1961.10.18.32-39 Paratype of *Haplochromis virginalis* (Kaduna form), coll. T.D.Iles, Nkhata Bay, Lake Nyasa 1 specimen 107.5 mm SL

D06E04 Chiweta Beach, Chilumba, 24-Feb-16, purchased from beach seiners, 1 specimen;

D12I04, I05, I08 Trawled from 30-40m depth SW of Makanjila, 2 March 2016, 3 specimens;

D14E01-3 trawled from 40m depth off Malembo, SW Arm, 4 March 2016, 3 specimens;

D17A01-D17A09, D17B03-D17B05, D17B07-B10 purchased from beach seiners, Palm Beach, SE Arm, 21 Jan 2017, 16 specimens;

D20E06-G02 purchased from Nkatcha fishermen, Lake Malombe, 24 Jan 2017, 17 specimens.

#### *Copadichromis virginalis* (Iles, 1960)

NHM 1961.10.18.31 Holotype of *Haplochromis virginalis* (Kaduna form), coll. T.D.Iles, Nkhata Bay, Lake Nyasa 92.4 mm SL

NHM 1961.10.18.32-39 Paratypes of *Haplochromis virginalis* (Kaduna form), coll. T.D.Iles, Nkhata Bay, Lake Nyasa 5 specimens 83.9-98.1 mm SL

NHM 1961.10.18.40-50 Paratypes of *Haplochromis virginalis* (Kajose form), coll. T.D.Iles, Nkhata Bay, Lake Nyasa 1 specimen 96.5 mm SL

D02G08-I01 purchased from fish traders, Nkhata Bay 21 Feb 2016, 14 specimens.

## Results

We first produced a neighbour-joining coalescent tree based on pairwise genetic differences at genome-wide SNP loci of the 51 recently sampled specimen. The tree showed relatively long terminal branches, consistent with the recent population separation and growth, but still resolved a sister relationship between two groups representing the Nkhata Bay specimens and all other ‘pure utaka’ specimens (Supplementary Figure 1). Block-bootstrap resampling in 100 kilobase windows confirmed 100% bootstrap support for the reciprocal monophyly of these groups. That said, only 3.5% of local gene trees supported this separation across all samples, an expected result given the observation that incomplete lineage sorting is widespread across Malawi cichlids (Malinsky et al. 2018).

All of the Nkhata Bay specimens collected freshly in 2016 were adult males with a very similar colour phenotype (Fig. 2c): dark grey body, pelvic, anal and caudal fins, with a bright yellow upper surface extending from the tip of the snout to the caudal fin. The dorsal fin was yellow with a narrow black stripe on the base, extending sharply upwards to the tip of the soft-rayed portion. The black stripe was sometimes bordered by white below the yellow area. Some of the paratypes of *H. virginalis* showed similar dark markings on the dorsal fin, particularly the 90.6mm SL, 86.9mm SL specimens, labelled as ‘Kaduna’ (Fig. 2b), as is the female holotype (Fig. 2a).

The other clade comprised both males and females. Male breeding colour was also dark grey to black with yellow on the upper surface, but the distribution of the yellow pigment was much more restricted, sometimes to the upper part of the dorsal fin, sometimes to the upperpart of the head and nape, but never extending far behind the first few dorsal fin rays and even then, only as a very thin band. The black in the dorsal fin always formed a much wider band at the base, sometimes extending right to the tip of the fin along its entire length (Fig. 2h, i). A wider dorsal fin black band was also shown by several of the *C. virginalis* types labelled as Kajose form, particularly the 100.7mm SL and 102mm SL specimens (fig. 2f). None of the *C. mloto* types showed any hint of male breeding colours (fig. 2e), similar to freshly collected females (fig. 2d): all were uniformly pale, slightly countershaded. Thus, male colours could not be used to characterise this species.

For better visualisation and to estimate species divergence time, we also computed a ‘split tree’ between samples where heterozygosity was subtracted from pairwise differences. The split tree yielded a qualitatively similar topology to the coalescent tree, again with 100% bootstrap support for the node separating the two clades (Fig. 3) with an estimated split time of the two clades at 53 kya (mutation rate based confidence interval 40-115 kya).

**Fig. 3.**
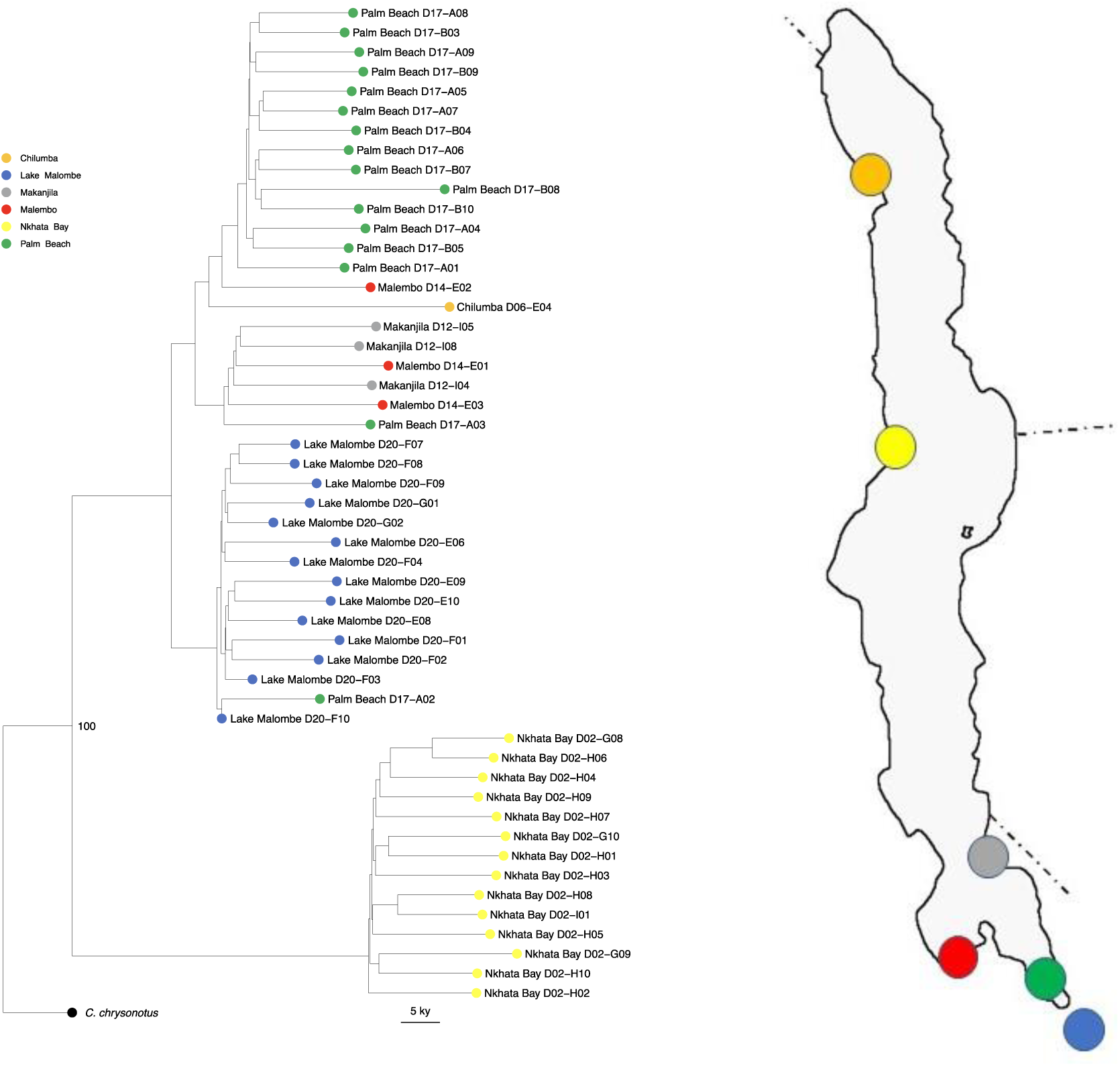
Neighbour joining ‘split tree’ based on 17 million SNPs from 51 whole genome sequences of *Copadichromis virginalis /mloto* specimens collected in 2016-17 from 6 localities in Lakes Malawi and Malombe. Code numbers refer to collection codes. Bootstrap support of 100% is shown for the divergence of *C. virginalis* and *C. mloto*.

Principal components analysis based on geometric morphometrics clearly separated the sequenced specimens into two groups: one comprised of the Nkhata Bay specimens which formed one major clade (Fig. 4, open orange triangles) and the other of all other specimens (Fig. 4, open green diamonds). The morphospace of the Nkhata Bay sequenced specimens was largely congruent with that of the types of *C. virginalis* assigned to the Kaduna morph, while the morphospace of the widely distributed clade largely encompassed both the types of *C. mloto* and the Kajose morph types of *C. virginalis*. Thus, we propose that the Kajose morph is in fact *C. mloto*.

**Fig. 4.**
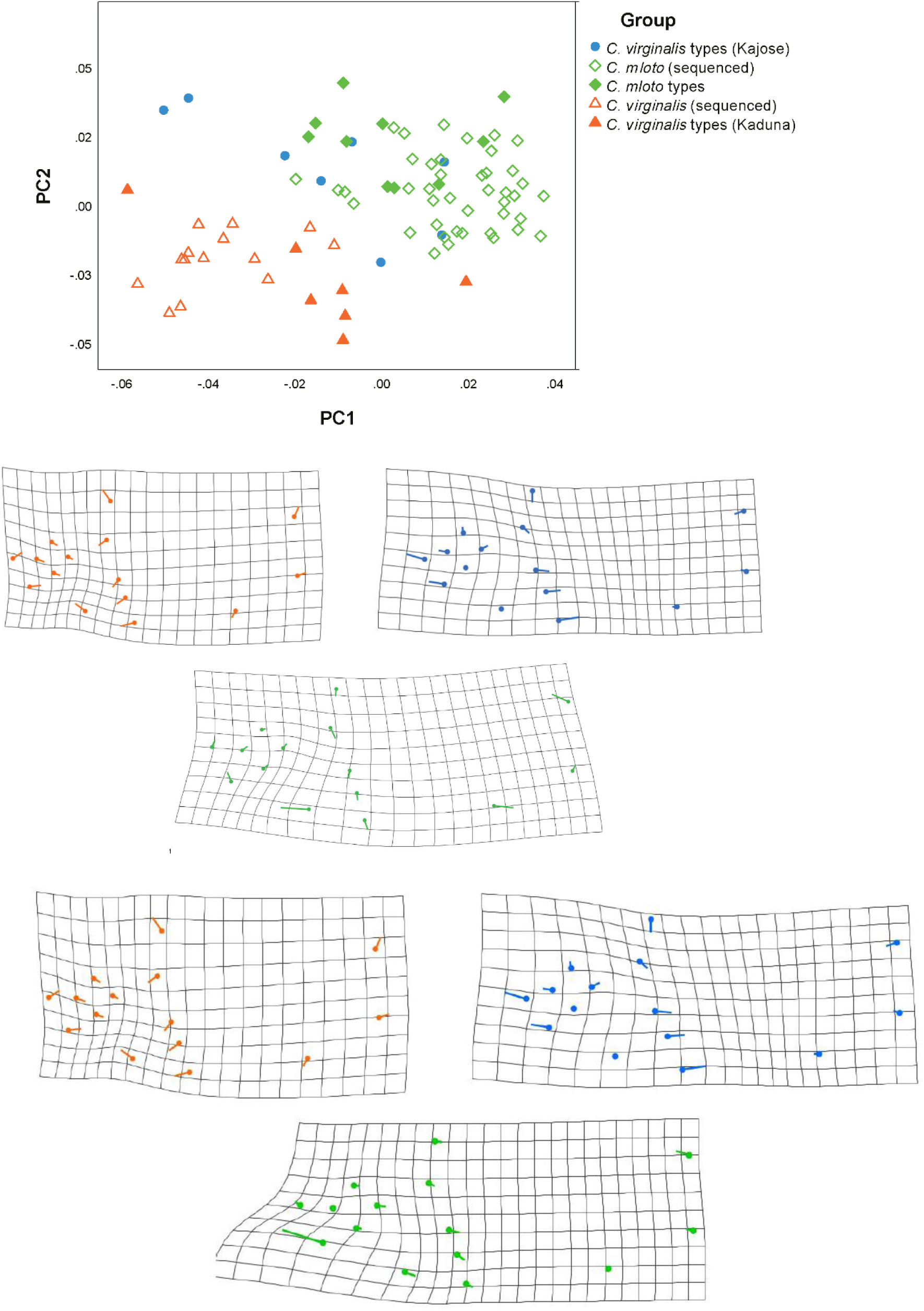
Above: Principal components analysis based on geometric morphometrics separates the *C. virginalis* specimens (orange, types shown by filled triangles) from the *C. mloto* (blue, types filled circles, types of *H. virginalis* ‘Kajose’ morph green filled diamonds); below: transformation grids for each group (same colour code). The points represent the consensus shape (mean shape of the group) and the line represents the shape change that corresponds to an increase of 0.1 units of Procrustes distance in the direction of the PC1.

**Fig. 5.**
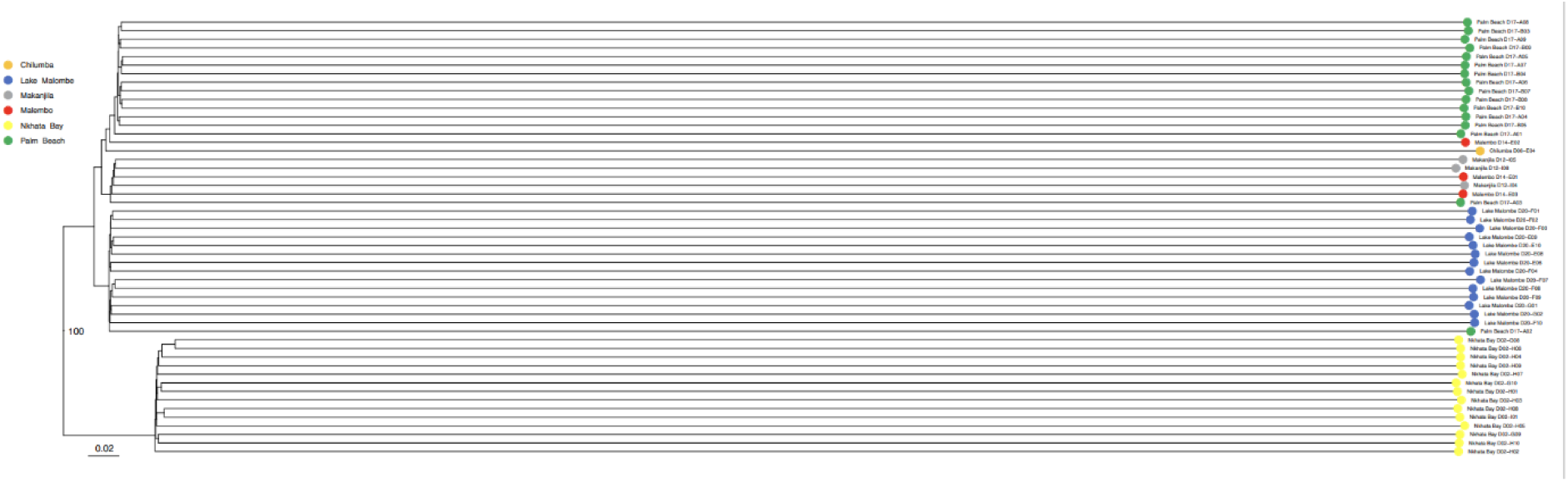
Phylogeny estimate using the Neighbour-Joining algorithm, with scale bar of 0.02 differences per kb accessible genome.

The first five principal components were responsible for 71% of the sample variance (see Supplementary table 1). Analysis of variance showed that all six comparisons between groups identified as *C. mloto* v *C. virginalis* were significantly different in body shape, including the Kaduna and Kajose (= *C. mloto*) types of *C. virginalis*, which were both comprised of a mix of sexes and which did not differ significantly in body (centroid) size.

Significant differences were also found in two of the four within-species comparisons. The difference between the Kaduna types of *C. virginalis* v the sequenced specimens in PC2 may have been the result of sexual dimorphism, as all the sequenced specimens were male, while the types were mixed sex. The difference between the sequenced *C. mloto* and the Kajose types may have been influenced by residual allometric effects as these two groups differed significantly in centroid size.

**Table 1.** Morphological comparisons among 5 populations: analysis of variance with post-hoc Tukey tests, with Bonferroni comparison for multiple testing, alpha <0.005 was taken as significant. CS= centroid size.

|  |  | *C. virginalis* |  | *C. mloto* |  |  |
| --- | --- | --- | --- | --- | --- | --- |
|  |  | Sequenced | Types (Kaduna) | Sequenced | Types (mloto) | Types (Kajose) |
| *C. virginalis* | Sequenced |  | PC2 | PC1 PC2 CS | PC1 PC2 | PC2 PC5 |
|  | Types (Kaduna) |  |  | PC1 PC2, CS | PC2 | PC2 |
| *C. mloto* | Sequenced |  |  |  | - | PC2 PC5, CS |
|  | Types (mloto) |  |  |  |  | - |
|  | Types (Kajose) |  |  |  |  |  |

## Discussion

The identification of *C. mloto* and *C*.*virginalis* has been confused since the original descriptions by Iles in 1960. Fryer &Iles (1972, p. 545) reported differences in the breeding seasons between the species, *C. virginalis* breeding May to June and *C. mloto* August to November, which makes it all the more puzzling that the original description contained no information on the breeding colour of *C. mloto* males, and no breeding males were included in the type series. They also reported *C. virginalis* and *C. mloto* (along with *Haplochromis kajose*!) were often found together in mixed species shoals (p. 295). Axelrod &Burgess (1986, but originally published in 1973 and periodically republished with similar/identical text but extra photos) showed a male and female *C. mloto* correctly identified and a female *C. virginalis* with very orange pelvic and anal fins, which may be *Mchenga flavimanus*. Fisheries data collection in the Monkey Bay research unit identified *C. mloto* as a common species in trawl catches in the southern arms of the lake (see Eccles &Trewavas, 1989), but this was later contradicted by Konings (1990), Turner (1996) and Snoeks &Hanssens (2004) who identified this species as *C. virginalis*.

Our combination of morphological investigation and sequencing of more recently collected material provides us with strong evidence that Iles’ Kajose form of *C. virginalis* is in fact conspecific with *C. mloto*, with the former representing deeper-bodied specimens including mature males and ripe females, while the types of the latter are slender individuals including spent fish. Notably, none of the *H. mloto* type material shows any sign of male breeding colours. Final certainty about the genetic identity of the type specimens relative to recent samples could only be obtained from their sequencing, which would be challenging given their preservation in formalin, but might become possible in the future (e.g., see Hahn et al., 2021). For the time being, the results presented here support the original assessment by Eccles &Trewavas (1989) that most pure utaka specimens commonly found in trawl catches correspond to *C. mloto* as do the species illustrated on Konings 1990 p. 256 (middle photo), Turner 1996 p. 53 and discussed on p. 67-69 as *C. virginalis*.

It is worth noting that the two male specimens shown by Turner (1996) show quite divergent breeding colours, as indeed do some of the specimens we collected, so it is quite possible that *C. mloto* as presently defined may represent a complex of species: resolving this would probably involve sampling males in breeding colour from Nkhata Bay – the type locality, but this has yet to be done. The specimens illustrated by Konings (2016, 324) as *C. mloto* do seem to correspond to our identification of that species. He reports males building bowers at depths of around 23m off Otter Point in the south of the lake, and describes the species as mainly being found over sand. Turner’s (1996) records and the sampling for the present study indicate the *C. mloto* is abundant in Lake Malombe and is found in Lake Malawi from the shallowest trawls (18m) down to 114m. We consider that the specimens examined by Anseeuw *et al*. (2008; 2011) as *C*. sp ‘virginalis kajose’ largely or entirely correspond to *C. mloto*: certainly those from Lake Malombe and SE Arm of Lake Malawi.

Konings (2016) also discusses *C. virginalis* and illustrates a number of males in breeding dress from a number of sites around the lake. These all correspond reasonably well to *C. virginalis* as we now define it, showing the characteristic narrow black dorsal fin base with the sharp uptick in the soft-rayed region. His *C. virginalis* are from Higga Reef (p.153), on the eastern shore of the lake across from Nkhata Bay on the West, and from Manda (p.154), also on the east coast but much further north. These males have a similar colour pattern to our Nkhata Bay specimens, but the yellow colour is replaced by pale blue: these would seem to represent geographic variants of *C. virginalis*, or allopatric sister species. His *C*. sp. ‘virginalis kajose’ from Kirondo (far north east) looks very similar to the Manda and Higga Reef specimens (same colour and dorsal fin markings) and does not correspond to *C. mloto*/kajose. This is also true of his *C*. sp. ‘firecrest mloto’ and *C. ilesi* both from Gome on the south eastern part of the lake near the Malawi/Mozambique border-their affinities would seem to lie with *C. virginalis* and not *C. mloto*. For this reason, we suggest the former should simply be referred to as *C*. sp. ‘firecrest’. In the original description of *C. ilesi*, it is apparent that Konings believed he was describing *C. virginalis* ‘kajose’ (which we now identify as *C. mloto*), although he did not include the Iles type specimens in his type series, but just specimens he had collected from Gome, illustrating males with a blue blaze. We agree with Snoeks and Hanssens (2004) that these are not *C. mloto/* ‘kajose’. In fact, photographs of *C. ilesi* from Gome in Konings (2016) look almost identical in colour to our *C. virginalis* (Kaduna) from Nkhata Bay, showing a male with a yellow blaze, and so perhaps *C. ilesi* might be best considered as a junior synonym of *C. virginalis*. However, not having examined the types of *C. ilesi* or sequenced specimens from Gome, we do not formally propose this.

The occurrence of two very differently coloured ‘*C. virginalis*’ forms at Gome – *C. ilesi* and *C*. ‘firecrest’ - suggests that some of these geographic colour forms are likely to deserve recognition as distinct species. The status of the populations illustrated by Konings (2016) as *C*. sp. ‘virginalis Chitande’ (NW coast around Chilumba) and *C*. sp. ‘virginalis gold’ (Nkanda, northeastern shore) is less clear. The former has pale blue upper parts and a weakly developed set of ‘virginalis’ black dorsal fin markings, while the latter has yellow upper parts, but no black in the dorsal at all. Irrespective of their status in relation to *C. virginalis*, we consider that none of these are referrable to *C. mloto*. Konings reports that these different variants of the *C. virginalis* complex all breed on rocky areas, constructing a bower consisting of a simple crater dug on a soft substrate underneath an overhanging rock, so that a complete circle is not produced, as the rock interrupts the circle. More certainty about the taxonomic status of different morphs will require more sampling and sequencing.

The estimated age of separation of the two species at around 50kya – albeit with substantial margin for error – is consistent with geological estimates for the last major refilling of the lake basin from around 100kya (see Scholz et al. 2007; Ivory et al. 2016). The strikingly long terminal branches observed in the NJ tree where within-individual hetrozygosity is retained (Supplementary Figure 1) reflect that cross-sample coalescence in many regions of the genome pre-dates the estimated splits, indicating substantial shared ancestral variation and incomplete lineage sorting. This accurately reflects the challenge in constructing robust phylogenetic estimates in Malawian cichlids, even with full genome sequence information (Scherz et al., 2022). A factor influencing this pattern may be a rapid population expansion around the time of speciation, perhaps co-incident with colonisation of deep-water habitats (Genner et al., 2010), while extensive introgression among species is probable and will further confound phylogeny estimation (Scherz et al., 2022).

## Conclusion

On the basis of our morphological and genomic studies, we conclude that *Copadichromis mloto* is an abundant species in a variety of fisheries in Lake Malawi, including the commercial trawls in the southern arms and the Nkatcha seines in Lake Malombe. It lives largely over sand/mud bottoms and can be recognised from its male breeding colours, body morphology and genome sequences. It includes the form previously recognised as Kajose from the type series of *Haplochromis virginalis*. The true *Copadichromis virginalis* is more rock-associated, building bowers up against overhanging rocks and characterised by breeding males with an extensive ‘blaze’ of bright colour (yellow at the type locality) on the whole of the upper surface and a distinctive black line on the lower part of the dorsal fin, showing a sharp uptick in the soft-rayed portion. Each of these two species may consist of a number of populations or geographic races with slightly different male colours, some of which may deserve recognition as further species.

## Acknowledgements

We are grateful to Mingliu Du, Richard Durbin, Milan Malinsky, Eric Miska, Mexford Mulumpwa, Bosco Rusuwa, Alexandra Tyers, Gregoire Vernaz, Richard Zatha for help with fieldwork, logistics and collecting permits and to Oliver Crimmen, James MacLaine and Simon Loader for help with accessing specimens in the Natural History Museum, London and to Richard Durbin for arranging for sequencing at the Wellcome Trust Sanger Institute. Funding was provided by a Wellcome Trust award (WT207492) to Richard Durbin and an Alborada Cambridge Africa Trust award to HS. AHvH was supported by FWO grand NR 11E0621N. HS was supported by the Flemish University Research Fund (BOF).

## Author contributions

Study was conceived and designed by Turner and Svardal, who also participated in the collection of material in Malawi. Specimens in the London Natural History Museum were examined and photographed by Turner. Analysis of sequence data and phylogeny estimation was performed and presented by Hooft van Huysduynen and Svardal. Geometric morphometric analysis was performed and presented by Crampton, who also prepared the images of the specimens. The first draft of the manuscript was written by Turner and all authors commented on previous versions of the manuscript. All authors read and approved the final manuscript.

## Appendix

**Table 2:** Summary of the Principal Components Analysis results, Geometric Morphometrics

| Variable | Eigenvalues<br>(x 10<sup>-4</sup>) | % Variance | Test among<br>groups;<br>Anova F<sub>4,76</sub>= | Correlation with<br>CS, Pearson's<br>r<sub>81</sub>= |
| --- | --- | --- | --- | --- |
| PC1 | 7.26 | 24.22 | 39.99; P<0.001 | -0.635; P<0.001 |
| PC2 | 5.73 | 19.11 | 28.99; P<0.001 | -0.265; P=0.017 |
| PC3 | 3.48 | 11.62 | 4.58; P=0.002 | -0.373; P=0.001 |
| PC4 | 2.95 | 9.83 | 3.68; P=0.009 | -0.122; P=0.028 |
| PC5 | 2.04 | 6.82 | 12.89; P<0.001 | -0.293; P=0.008 |
| Centroid<br>Size | n/a | n/a | 26.59; P<0.001 | n/a |

## References

Anseeuw, D., G. E. Maes, P. Busselen, D. Knapen, J. Snoeks, & E. Verheyen, 2008. Subtle population structure and male-biased dispersal in two Copadichromis species (Teleostei, Cichlidae) from Lake Malawi, East Africa. Hydrobiologia 615: 69–79.

Anseeuw, D., J. A. Raeymaekers, P. Busselen, E. Verheyen & J. Snoeks. 2011. Low genetic and morphometric intraspecific divergence in peripheral Copadichromis populations (Perciformes: Cichlidae) in the lake Malawi basin. International Journal of Evolutionary Biology 2011: 835946, 11 pp.

Axelrod, H. R. & W. E. Burgess, 1986. African Cichlids of Lakes Malawi and Tanganyika. 11th ed. T.F.H. Publications, Neptune City, NJ.

Bookstein, F. L. 1989. Principal Warps: thin-plate splines and the decomposition of deformations. IEEE Trans. Pattern Anal. Mach. Intell. 11: 567–585.

Danecek, P., J.K. Bonfield, J. Liddle, J. Marshall, V. Ohan, M.O. Pollard, A. Whitwham, T. Keane, S.A. McCarthy, R.M. Davies & H. Li. Twelve years of SAMtools and BCFtools. Gigascience 2021:10(2), giab008.

Eccles, D. H., & E. Trewavas, 1989. Malawian cichlid fishes. The classification of some Haplochromine genera. Lake Fish Movies, Herten, Germany, 335 pp.

Fryer, G. & T. D. Iles, 1972. The Cichlid Fishes of the Great Lakes of Africa. Oliver & Boyd, Edinburgh, 641 pp.

Genner, M. J., M. E. Knight, M. P. Haesler & G. F. Turner, 2010. Establishment and expansion of Lake Malawi rock fish populations after a dramatic Late Pleistocene lake level rise. Molecular Ecology. 19:170–182.

Hahn, E. E., M. R. Alexander, A. Grealy, J. Stiller, D. M. Gardiner & J. Stiller. 2021 Unlocking inaccessible historical genomes preserved in formalin. Molecular Ecology Resources (online ahead of publication)

Iles, T. D. 1960. A group of zooplankton feeders of the genus Haplochromis (Cichlidae) in Lake Nyasa. Annals and Magazine of Natural History (13) 2, 1959: 257–280.

Klingenberg, C. P. 2011. MorphoJ: An Integrated Software Package for Geometric Morphometrics. Molecular Ecology Resources. 11: 353–357.

Konings, A. 1990. Cichlids and All the Other Fishes of Lake Malawi. T.F.H. Publications, Neptune City, NJ.

Konings, A. 1999. Descriptions of three new Copadichromis species (Labroidei; Cichlidae) from Lake Malawi, Africa. Tropical Fish Hobbyist 47 (9): 62–84.

Konings, A. 2016. Lake Malawi Cichlids in their Natural Habitat. 5th Edn. Cichlid Press, El Paso TX.

Malinsky, M., H. Svardal, A.M. Tyers, E.A. Miska, M.J. Genner, G.F. Turner & R. Durbin, 2018. Whole genome sequences of Malawi cichlids reveal multiple radiations interconnected by gene flow. Nature Ecology & Evolution 2: 1940–1955.

Parés-Casanova, P. M., A. Salamanca-Carreño, R. A. Crosby-Granados & J. Bentez-Molano. 2020. A Comparison of Traditional and Geometric Morphometric Techniques for the Study of Basicranial Morphology in Horses: A Case Study of the Araucanian Horse from Colombia. Animals. 10: 118.

Rohlf, F. J. 2004. TpsDig Version 1.4. Department of Ecology and Evolution. State University of New York at Stony Brook, New York.

Rohlf, F. J. 2015. The Tps series of software. Hystrix, the Italian Journal of Mammalogy. 26: 9–12

Rohlf, F. J. & D. Slice. 1990. Extensions of the Procrustes Method for the Optimal Superimposition of Landmarks. Systematic Zoology. 39: 40–59.

Rusinko, J. & M. McPartlon, 2017. Species tree estimation using Neighbor Joining. Journal of theoretical biology: 414, 5–7.

Scherz, M. D., P. Masonick, A. Meyer & C. D. Hulsey 2022. Between a rock and a hard polytomy: phylogenomics of the rock-dwelling cichlids of Lake Malawi. Systematic Biology (online ahead of publication).

Scholz, C. A., T. C. Johnson, A. S. Cohen, J. W. King, J. A. Peck, J. T. Overpeck, M. R. Talbot, E. T. Brown, L. Kalindekafe, P. Y O. Amoaka, R. P. Lyons, T. M. Shanahan, I. S. Castañeda, C. W. Heil, S L. Forman, L. R. McHargue, K. R. Beuning, J. Gomez, J. Pierson (2007) East African megadroughts between 135 and 75 thousand years ago and bearing on early-modern human origins. Proceedings of the National Academy of Sciences USA. 104: 16416–16421.

Snoeks, J. 2004. Introduction. In Snoeks, J. (ed) The Cichlid Diversity of Lake Malawi/Nyasa/Niassa: Identification, Distribution and Taxonomy. Cichlid Press, El Paso, TX: 9–11.

Snoeks, J. & M. Hanssens, 2004. Identification guidelines to other non-mbuna. In Snoeks, J. (ed) The Cichlid Diversity of Lake Malawi/Nyasa/Niassa: Identification, Distribution and Taxonomy. Cichlid Press, El Paso, TX: 266–310.

Svardal, H., F. X. Quah, M. Malinsky, B. P, Ngatunga, E. A. Miska, W. Salzburger, M. J Genner, G. F Turner, R. Durbin, 2020. Ancestral hybridization facilitated species diversification in the Lake Malawi cichlid fish adaptive radiation, Molecular Biology and Evolution 37: 1100–1113.

Turner, G. F. 1996. Offshore Cichlids of Lake Malawi. Cichlid Press, Lauenau. 240 pp.

